# A drug discovery-oriented non-invasive protocol for protein crystal cryoprotection by dehydration, with application for crystallization screening

**DOI:** 10.1101/2021.11.23.469721

**Authors:** Dom Bellini

**Affiliations:** MRC Laboratory of Molecular Biology, Francis Crick Avenue, Cambridge Biomedical Campus, Cambridge CB2 0QH, United Kingdom

## Abstract

In X-ray macromolecular crystallography, cryoprotection of crystals mounted on harvesting loops is achieved when the water in the sample solvent transitions to vitreous ice before crystalline ice forms. This is achieved by rapid cooling in liquid nitrogen or propane. Protocols for protein crystal cryoprotection are based on either increasing environmental pressure or reducing the water fraction in the solvent. This study presents a new protocol for cryoprotecting crystals. It is based on vapour diffusion dehydration of the crystal drop to reduce the water fraction in the solvent by adding a highly concentrated salt solution, 13 M potassium formate (KF13), directly to the reservoir. Cryoprotection by the KF13 protocol is non-invasive to the crystal, high throughput, not labour intensive, can benefit diffraction resolution and ligand binding, and is very useful in cases with high redundancy such as drug discovery projects which utilize very large compound or fragment libraries. Moreover, an application of KF13 to discover new crystal hits from clear drops of equilibrated crystallization screening plates is also shown.

## 1. Introduction

In macromolecular X-ray crystallography, it is important to perform data collections at cryogenic temperatures (usually around 100 K) where crystal radiation damage is significantly slower, especially at high intensity synchrotron radiation sources (Kmetko *et al*., 2006, Owen *et al*., 2006). Cryoprotection of loop-harvested crystals is intended to achieve water transition to vitreous ice before crystalline ice formation, upon flash cooling in liquid nitrogen. Crystalline ice formation should be avoided. First, it compromises diffraction quality by destabilizing the crystal lattice due to its volume expansion compared to liquid water (Haas & Rossmann, 1970, Juers & Matthews, 2001, 2004, Kriminski *et al*., 2002, Low *et al*., 1966). These effects cause crystal disorder and/or non-isomorphism. Second, it causes large variations in background counts of diffraction images due to X-ray diffraction by cubic and hexagonal ice at specific Bragg angles (the so-called “ice rings”) (Burkhardt *et al*., 2012, Fuentes-Landete *et al*., 2015, Parkhurst *et al*., 2017, Thorn *et al*., 2017).

There are two main strategies to cryoprotect macromolecular crystals, which aim to either 1) increase the environmental pressure (Burkhardt *et al*., 2012, Kim *et al*., 2005, Thomanek *et al*., 1973) or 2) reduce the solvent fraction to below the glass-transition phase of water (Pflugrath, 2015). The latter, reduction of solvent fraction, can be achieved by either a) soaking crystals in cryosolutions enriched with cryoprotective agents such as sugars, salts, polyethylene glycols, glycerol, and various others (Bujacz *et al*., 2010, Gulick *et al*., 2002, Holyoak *et al*., 2003, Hope, 1988, Marshall *et al*., 2012, Pemberton *et al*., 2012, Rubinson *et al*., 2000, Vera & Stura, 2014) or b) dehydration. While in some cases an adequate cryoprotective agent is already present in the crystallization buffer, very often an additional step needs to be performed where the crystal is transferred to a cryoprotected solution; this process can be laborious and damaging to crystals due to handling and osmotic stress, respectively. Crystal handling is avoided in procedures that make use of acoustic nanodroplet ejectors (e.g., the Echo acoustic liquid handler from Labcyte inc.) for precise placement of cryoprotective agents directly within the crystallization drop but away from the crystals and towards the drop edges to allow gradual gentle diffusion; however, the set-up of such a pipeline is neither straightforward nor inexpensive, including also the requirement for crystal plate imaging facilities (Collins *et al*., 2017). In addition, a low throughput but efficient protocol has been reported for crystal cryoprotection by using vapour diffusion of volatile alcohols (Farley & Juers, 2014). Dehydration studies of macromolecular crystals instead have usually been aimed at improving data resolution rather than cryoprotection (Abergel, 2004, Esnouf *et al*., 1998, Heras *et al*., 2003, Kiefersauer *et al*., 2000). However, two of these studies have also reported that some crystals, dehydrated either using a humidity control device (Sanchez-Weatherby *et al*., 2009) or replacing the reservoir in the crystallization plate with NaCl solutions (Douangamath *et al*., 2013), no longer required being soaked in cryosolutions to prevent crystalline ice formation during flash cooling.

In this study, a new cryoprotection protocol is described in which a solution of 13 M potassium formate (KF13) is added in a single step directly to the plate reservoir. This dehydrates the crystal drop overnight by vapour diffusion, thereby cryoprotecting the crystal. This method successfully cryoprotected six different crystal systems, which were grown in conditions containing different salts and polyethylene glycols. The amount of KF13 added to achieve cryoprotection, without over-dehydrating the crystal at the expense of diffraction quality, depends on both the components of the crystallization solution and crystal solvent content. It was shown to vary between 4-20% of the final volume (reservoir plus KF13).

Moreover, this work also showed that adding KF13 to the reservoir of previously equilibrated crystallization screening plates can promote, through further dehydration by vapour diffusion, the formation of new crystals from “idled” clear drops. This approach provides a new high throughput protocol to recycle unsuccessful crystallization screening conditions. Clear drops usually accounts for around 50% of all conditions in a crystallization screening experiment.

## 2. Materials and Methods

### 2.1. Chemicals and protein crystals

Precipitants, such as polyethylene glycols and salts, buffers and chemicals to prepare dehydrating solutions such as potassium formate were purchased from Sigma. The commercially available proteins *Thaumatococcus daniellii* (an african plant) thaumatin, *Gallus gallus* (hen egg white) lysozyme and *Canavalia ensiformis* (jack bean) concanavalin A were also purchased from Sigma. Crystals of *Staphylococcus aureus* FtsA filaments, *Homo sapiens* hetero-pentamer Cenp-OPQUR complex and glutamate receptor ligand binding domain in complex with agonist (GluLBD) were kindly donated by LMB researchers Danguole Ciziene, Stan Yatskevich and Christina Heroven, respectively. Crystallization sitting drops were prepared on the Mosquito nanoliter liquid handler (STP Labtech) at 293 K by dispensing equal volumes (200 nL) of protein and reservoir solutions. For the study on cryoprotection by dehydration, crystals of lysozyme, concanavalin A and thaumatin were prepared at various precipitant concentrations, to study both direct effects of precipitant concentration on cryoprotection, and also to produce crystals of different sizes, since crystal size can also affect cryoprotection. To differentiate between the same protein samples that were crystallized at different precipitant concentrations, the following acronyms were used throughout this manuscript:

a) lyso07, lyso08 and lyso12 describe lysozyme crystals grown in 0.7, 0.8 and 1.2 M NaCl, respectively; b) conc11, conc13 and conc14 represent concanavalin A crystals grown in 11, 13 and 14% PEG 6k, respectively; c) thau06 and thau10 describe thaumatin crystals grown in 0.6 and 1.0 M NaK tartrate, respectively. The detailed list of crystallization conditions is presented in Tables 1 and S1, respectively, for the study on cryoprotection by dehydration (Table 1) and promotion of crystal nucleation in already vapour diffusion equilibrated crystal drops (Table S1).

**Table 1.** Crystal sizes and forms, crystallisation conditions and X-ray beam size used in the study of cryoprotection by the KF13 protocol.

| Sample | Protein concentration (mg/ml) | Crystallization conditions | Crystal size ($\mu\text{m}$) and shape | X-ray beam size ($\mu\text{m}$) | Space group |
| --- | --- | --- | --- | --- | --- |
| FtsA | 14 | 8% PEG 8k, 8% PEG 1k, 200 mM Li<sub>2</sub>SO<sub>4</sub> and tris pH 8.5 | $\approx 300$ (rods) | 50 x 50 | I222 |
| Cenp-OPQUR | 15 | 15% PEG 2k mme, 40 mM NaFormate and 200 mM bis-tris propane pH 6.9 | $\approx 400$ (thin plates) | 50 x 50 | C2 |
| GluLBD | 10 | 20% PEG 4k, 200 mM CaCl<sub>2</sub> and 100 mM Tris pH 8 | $\approx 150$ (rods) | 400 x 400 (home source) | P2<sub>1</sub>2<sub>1</sub>2<sub>1</sub> |
| Lyso07 | 12 | 0.7 M NaCl and 50 mM sodium acetate pH 4.5 | $\approx 50$ (diamonds) | 11 x 5 | P4<sub>3</sub>2<sub>1</sub>2 |
| Lyso08 | 40 | 0.8 M NaCl and 50 mM sodium acetate pH 4.5 | $\approx 50$ (diamonds) | 11 x 5 | P4<sub>3</sub>2<sub>1</sub>2 |
| Lyso12 | 40 | 1.2 M NaCl and 50 mM sodium acetate pH 4.5 | $\approx 200$ (diamonds) | 11 x 5 | P4<sub>3</sub>2<sub>1</sub>2 |
| Conc11 | 12 | 11% PEG 6k, 5% pentanediol and 100 mM Hepes pH 7.5 | $\approx 200$ (cuboids) | 11 x 5 | I222 |
| Conc13 | 12 | 13% PEG 6k, 5% pentanediol and 100 mM Hepes pH 7.5 | $\approx 200$ (cuboids) | 11 x 5 | I222 |
| Conc14 | 12 | 14% PEG 6k, 5% pentanediol and 100 mM Hepes pH 7.5 | $\approx 200$ (cuboids) | 11 x 5 | |
| Thau06 | 25 | 0.6 M NaK tartrate and 100 mM potassium phosphate pH 6.8 | $\approx 200$ (diamonds) | 11 x 5 | P4<sub>1</sub>2<sub>1</sub>2 |
| Thau10 | 25 | 1.0 M NaK tartrate and 100 mM potassium phosphate pH 6.8 | $\approx 50$ (diamonds) | 11 x 5 | P4<sub>1</sub>2<sub>1</sub>2 |

### 2.2. Dehydration of crystallization drops

Protein crystallization was carried out in 96-well MRC plates (SWISS-CI) containing 80 μl of reservoir solutions. This type of plate accommodates up to 100 μl of reservoir. Required volumes of KF13, ranging from 0-20 μl, were added directly to the reservoir by either a) removing completely the sealing tape and using a multi-channel pipette before quickly re-sealing the whole plate with 3-inch wide Crystal Clear sealing tape (Hampton Research), in case of high throughput experiments such as the study of promoting crystal nucleation in already equilibrated clear drops, or b) making a small cross incision in the sealing tape with a blade and using a single-channel pipette, before re-sealing the cut with a small piece of crystal clear tape (Hampton), in the case of low throughput experiments such as the study of cryoprotection by dehydration. For crystals of FtsA, Cenp-OPQUR and GluLBD, due to the limited amounts of samples, experiments were performed only once. In contrast, with the commercially available samples, experiments were performed in duplicates (thaumatin) or triplicates (lysozyme and concanavalin A); however, the concentration of precipitants used in crystallization was slightly varied amongst replicate plates to produce crystals of different sizes (see previous section and Table 1).

### 2.3. Diffraction data collection

Single diffraction images were acquired on beamline I24 (Diamond Light Source) and the home source (FrE+ superbright, Rigaku) for crystals of Cenp-OPQUR and GluLBD, respectively. Complete datasets instead were collected for crystals of FtsA on beamline I24 and of lysozyme, concanavalin A and thaumatin on I04 (Diamond Light Source). All diffraction data were collected at 100 K and autoprocessed with Xia2 DIALS (Winter *et al*., 2018).

## 3. Results and Discussion

### 3.1. KF13 is an ideal solution for crystal drop dehydration by vapour diffusion

All crystallization plates used in this study are 96-well MRC plates. These allow a maximum reservoir volume of 100 μl. Ready-to-use crystallization screening plates prepared at the LMB crystallization facility contain 80 μl of reservoir in each well (significantly lower volumes are not ideal for storing at 10° C due to evaporation issues) allowing addition of a maximum of 20 μl of extra solution. To achieve further dehydration of equilibrated crystal drops via vapour diffusion by adding some 0-20 μl solution to the reservoir, the overall osmotic strength of 80 μl of reservoir plus x μl of very concentrated solution (where x equals 0-20) must be stronger than that of the original 80 μl reservoir (since the osmotic strength of an equilibrated crystal drop is approximately the same as that of the reservoir). Therefore, to counteract the diluting effect that adding extra solution has on the original reservoir and prevent rehydration rather than dehydration, the osmotic strength of the added solution must be very high (since the reservoir already contains high concentrations of precipitants). A schematic drawing of this process is showed in Fig. 1.

**Figure 1.**
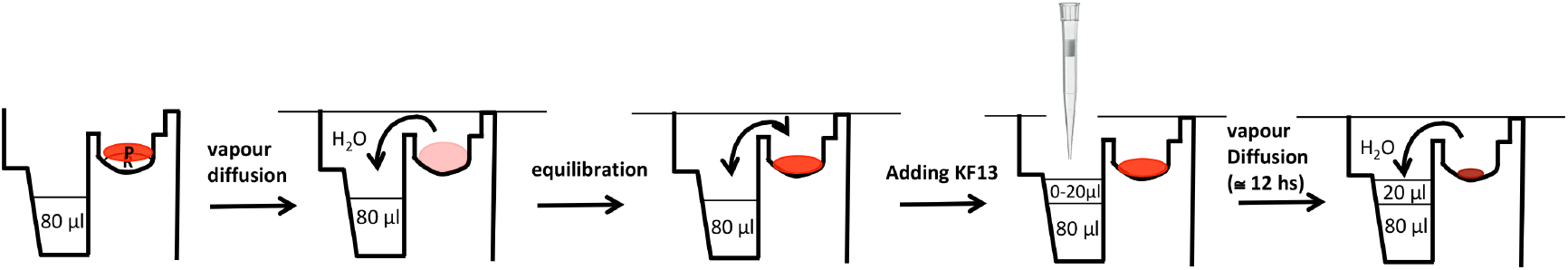
Schematic drawing of the procedures involved in crystal drop dehydration using the KF13 protocol.

A number of potential candidates to generate very highly concentrated solutions with high osmotic strength were tested (Table 2). Interestingly, it was not possible to achieve the reported maximum solubility at room temperature of most of the tested substances. Thus the tests were carried out at the maximum achievable solubility without heating solutions to reach higher concentrations (this way the solution is more stable and can be store indefinitely at room temperature without the problem of the solute precipitating in case of a reduction in the temperature) (Table 2). Solutions of caesium acetate and fructose were excluded due to their expense and being too viscous for pipetting, respectively (Table 2). NaCl, K-acetate, CsCl and Cs-phosphate failed to dehydrate a large number of crystallization drops whose reservoirs contained a significant amount of precipitant (e.g., more than 1.5 M ammonium sulphate or more than 20% of PEG with molecular weight higher than 1000 da), whereas KNO_2_ was excluded because it reacted with solutions containing (NH_4_)_2_SO_4_, releasing nitrogen gas [2KNO_2_ + (NH_4_)_2_SO4 → 2N_2_ + K_2_SO_4_ + 4H_2_O], as judged by the bubbling reservoir, which caused the sealing tape to open up in some places due to the building up of pressure in the well (Table 2). Solutions of 10 M Li-iodide and 13 M K-formate (KF13) proved to be equally efficient in causing crystal drop dehydration in all crystallization conditions tested (Table 2). However, KF13 was chosen as the preferred solution because it is ten times less expensive than Li-iodide. Figure 2 shows how adding 20 μl of KF13 to different 80 μl of reservoirs containing different precipitants at different concentrations significantly dehydrated all crystal drops.

**Table 2.** Chemicals tested for the preparation of suitable solutions to be added to crystallisation plate reservoirs for drop dehydration by vapour diffusion.

| Chemical | Theoretical max concentration (M) | Achieved max concentration (M) | Performance in dehydration experiments |
| --- | --- | --- | --- |
| NaCl | 6 | 6 | too weak |
| Fructose | 20 | 8 | too viscous |
| K-acetate | 20 | 8 | too weak |
| CsCl | 8 | 6 | too weak |
| Cs-acetate | 40 | 14 | too expensive |
| Cs-phosphate | 10 | 8 | too weak |
| KNO<sub>2</sub> | 25 | 10 | Formation of gas in combination with AmSO<sub>4</sub> |
| Li-iodide | 10 | 10 | OK |
| K-formate | 30 | 13 | OK |

**Figure 2.**
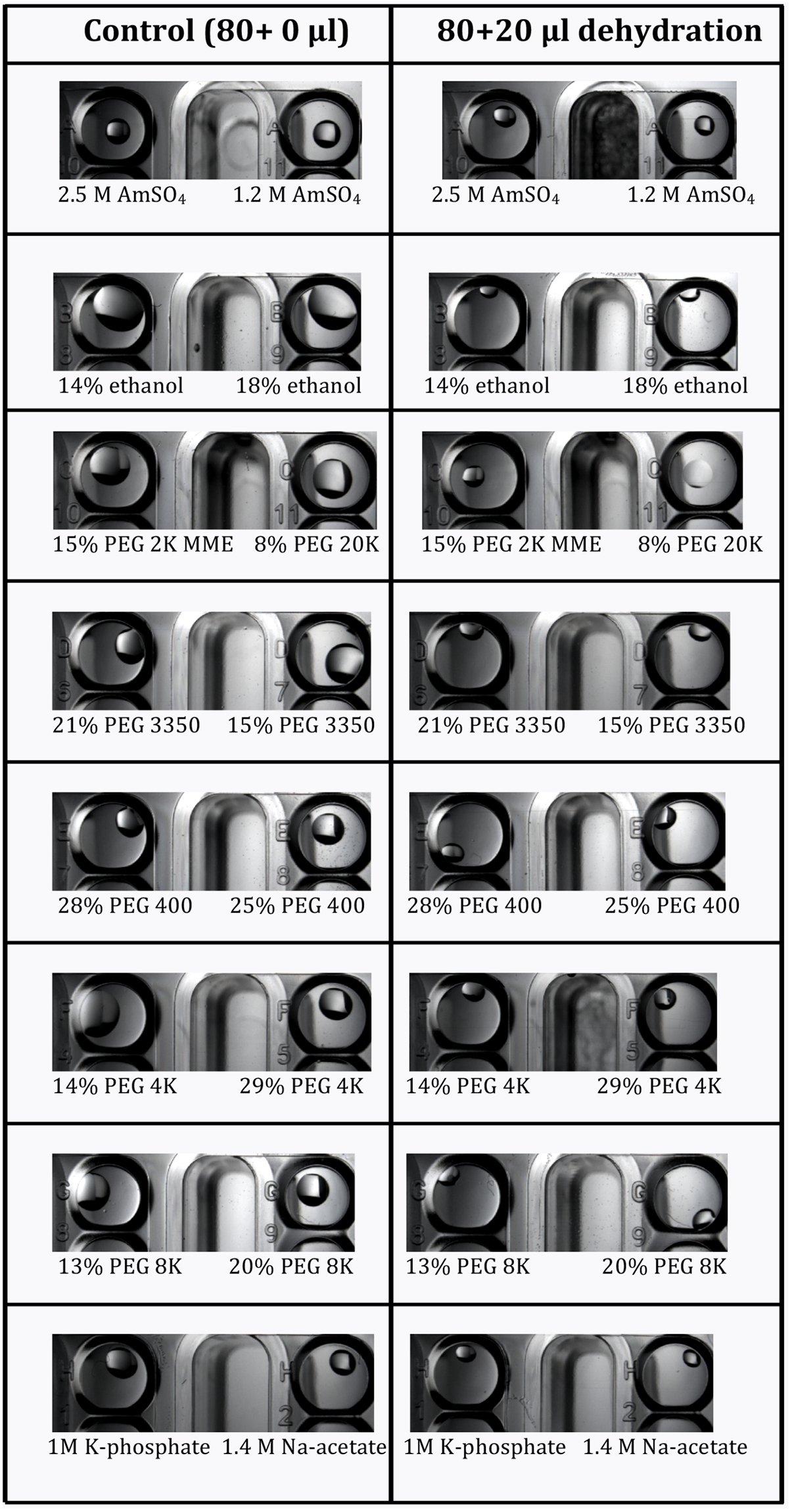
Dehydrating effects on crystallization drops from significantly different conditions after 24 hours from adding KF13 to the reservoirs.

### 3.2. Cryoprotection by KF13 dehydration

Tests were conducted using six different protein samples, namely FtsA, Cenp-OPQUR, GluLBD, lysozyme, concanavalin A and thaumatin (Table 1). Moreover, crystals of different sizes were produced for lysozyme, concanavalin A and thaumatin by varying precipitant concentrations (Table 1 and Fig. 1S). Crystal drops were dehydrated by adding 2, 4, 6, 8, 10, 15 or 20 μl of KF13 directly to the reservoirs (80 μl) and left to equilibrate for 12-24 hours. Crystals were then harvested in nylon loops and flash-frozen in liquid nitrogen. Diffraction images highlighting the transition from ice to glass as judged by the disappearance of ice rings relative to the amounts of KF13 added to the reservoirs are shown in Fig. S2-S7 for each of the six crystals forms, respectively. Table 3 summarizes these results with an approximate value for the minimum amount of KF13 needed to achieve cryoprotection in relation to the type and amount of precipitant, and the solvent content of the crystals. Despite the solvent content as high as 72%, crystals from drops containing polyethylene glycols (ranging from 8-20%) only required 4-10 μl of KF13 to achieve cryoprotection, whereas in the case of drops containing salts as precipitants, the amounts of KF13 required was markedly higher, being in the range of 15-20 μl, despite solvent content as low as 32%. As shown in Table 3, the variation in the size of lysozyme and thaumatin crystals (Table 1 and Fig. S1) did not appear to have any effects on cryoprotection. Most important is the fact that this protocol has achieved cryoprotection even in cases where the precipitant is a very poor cryoprotectant such as NaCl (Berejnov *et al*., 2006), suggesting that this KF13-based approach can be applied to crystals grown in most if not all crystallization conditions currently in use in macromolecular crystallography. In this study the diameters of the harvesting nylon loops were chosen to approximately match the size of the crystal, and the X-ray beam cross section was smaller than the crystal size (Fig. S1).

**Table 3.** Summary of KF13 volumes required to achieve cryoprotection by vapour diffusion dehydration in crystals grown from different conditions and with different solvent contents.

| Sample | precipitant | Solvent content (%) | Minimum volume of KF13 required for cryoprotection ($\mu\text{l}$) |
| --- | --- | --- | --- |
| FtsA | 8% PEG 8k, 8% PEG 1k, | 72 | 10 |
| GluLBD | 20% PEG 4k | 57 | 4 |
| Cenp-OPQUR | 15% PEG 2k mme | 50 | 8 |
| Lyso_07 | 0.7 M NaCl | 32 | 20 |
| Lyso_08 | 0.8 M NaCl | 32 | 20 |
| Lyso_12 | 1.2 M NaCl | 32 | 20 |
| Conc_11 | 11% PEG 6k | 46 | 6 |
| Conc_13 | 13% PEG 6k | 46 | 6 |
| Conc_14 | 14% PEG 6k | 46 | 5 |
| Thau_06 | 0.6 M NaK tartrate | 55 | 15 |
| Thau_10 | 1.0 M NaK tartrate | 55 | 15 |

Complete datasets were collected for FtsA, lysozyme, concanavalin A and thaumatin crystals. Diffraction data processing showed in all cases a strong correlation between disappearance of ice rings, lower mosaicity and higher resolution (Fig. 3). Continuing to increase the amount of KF13 passing the point of ice ring disappearance eventually caused an increased mosaicity and reduced resolution (except in the case of lysozyme crystals where cryoprotection was only reached when adding the maximum amount of KF13 (20 μl) allowed by the well size without the need to remove any of the 80 μl of reservoir); this also appeared to be true in the case of single-image-collections of GluLBD and Cenp-OPQUR samples as judged from the deterioration in resolution in the diffraction images of crystals from drops dehydrated with KF13 volumes above the optimal 4 and 10 μl, respectively (Fig. S3 and Fig. S4). The overall amounts of KF13 that can be added for achieving optimal cryoprotection without degrading the diffraction resolution can vary significantly depending on the sample, ranging from a narrow 8-10 μl interval for FtsA crystals to a wider 5-15 μl for concanavalin A crystals (Fig. 3).

**Figure 3.**
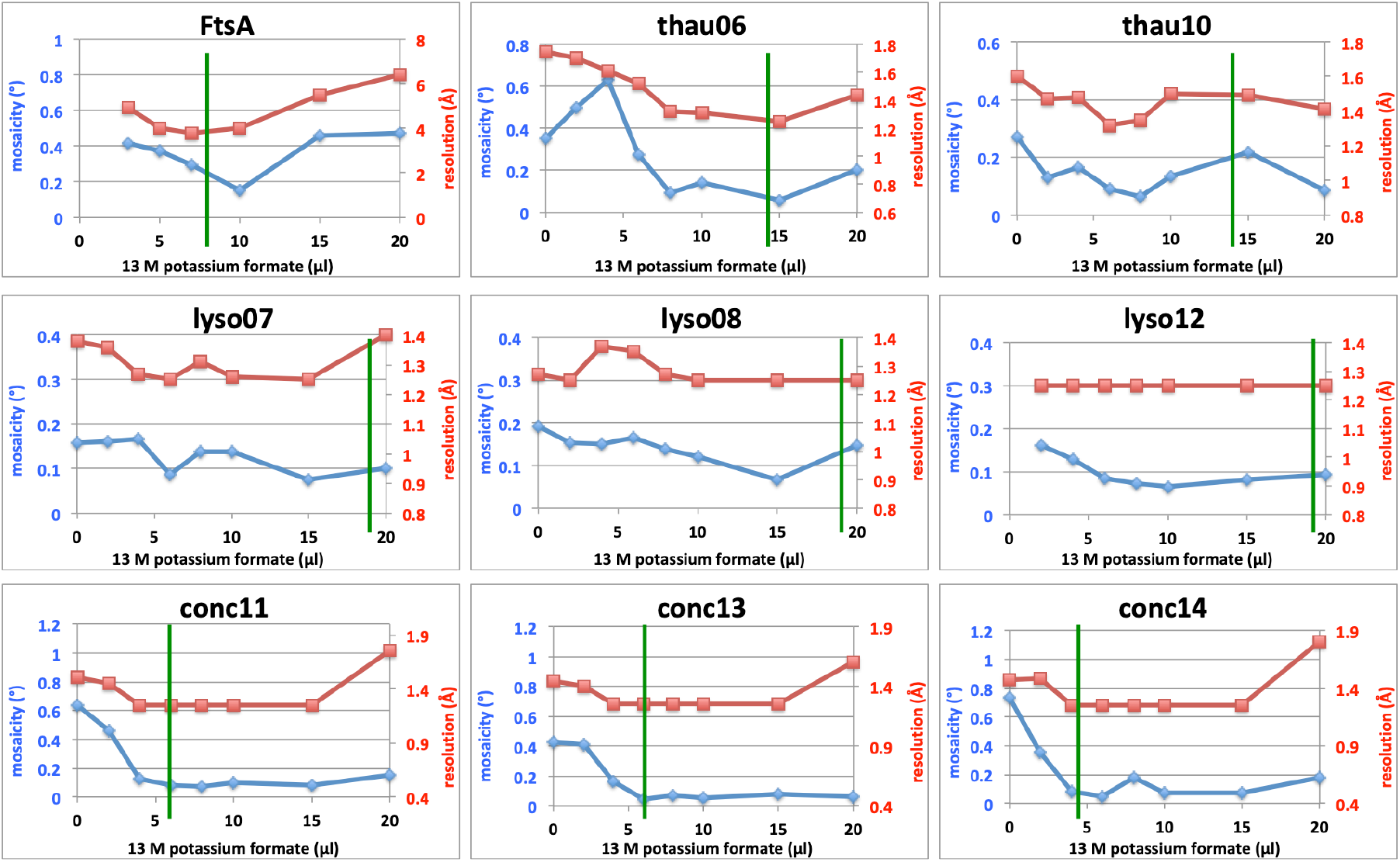
Correlation between amounts of KF13 used for crystal drop dehydration and both mosaicity and diffraction resolution of datasets collected for different crystal samples. The green line indicates the minimum value of KF13 volume that caused the ice rings to disappear.

As expected, crystal drop dehydration induces a hypertonic stress to the unit cell of the crystals causing it to shrink proportionally to the amount of KF13 added to the reservoir, except in the case of thaumatin where no contraction of the unit cell was observed, perhaps suggesting a very rigid lattice (Fig. S8). All other samples underwent an anisotropic contraction of the unit cell, except concanavalin A crystals, where all three unit cell axes shrunk (Fig. S8). As reported in the Introduction, unit cell shrinkage and consequently reduction in solvent content of crystals by dehydration has been exploited in a number of cases to improve lattice order and consequently diffraction resolution, especially with crystals of membrane proteins crystallized with detergents. Therefore, cryoprotection by KF13 dehydration may also lead to improved crystal order and better diffraction resolution compared to other cryoprotection techniques that do not exert hyperosmotic pressure on the crystal. Since lattice shrinkage is shown to be proportional to the added volume of KF13, the overall amount of KF13 required to achieve full cryoprotection (e.g., 6 μl) could be added gradually (e.g., 2 μl every 24 hours) thereby minimizing hypertonic shock and optimizing diffraction resolution in certain cases.

The KF13 cryoprotection method is ideal for crystallographic drug discovery projects for a number of reasons. 1) In standard cryoprotection protocols the crystal is transferred to a cryoprotective agent-containing drop, which ideally should also contain the ligand of interest; however, it can be very labour intensive to prepare all the different cryoprotectant solutions each containing a unique ligand (or even cocktails of ligands) from a large library for each different ligand-complex crystal. This is unnecessary with the KF13 method since the crystallization solution remains untouched. 2) Since KF13 cryoprotection is achieved by dehydration, this means that the concentration of ligand in the drop will rise causing binding occupancies to increase, thus improving the probability to observe the ligand in the electron density map. This latter consideration is also true in the case of *ab initio* structure determination experiments involving metal atom soaks. 3) While the KF13 method works well in low throughput experiments, at the same time it is also ideal for very large projects such as screening of large libraries of compounds or fragments. In these cases, high throughput can be achieved by either adding KF13 manually using multi-channel pipettes or via an automated pipeline using liquid handling robots.

The crystal drop dehydration method of replacing the reservoir with a NaCl solution from Douangamath *et al*. (2013) was developed with the aim to rapidly screen effects of dehydration on the diffraction resolution of crystals *in situ*. The KF13 protocol is also perfectly suited for such a purpose, with the advantage that KF13 is added in a single step without necessitating prior removal of the reservoir.

### 3.3. KF13 can aid crystal hit discovery by dehydrating clear drops in pre-equilibrated screening plates

Standard protocols for identification of new crystal hits consist of screening over one to two thousand different crystallization conditions using normally 96-well plates. An important parameter in this process is the choice of protein concentration, which, as a rule of thumb, is taken as the concentration that produces around 50% of clear drops (intended as no observable protein precipitation) immediately after setting up the plates. This means that in any given crystal hit screening experiment, even after achieving drop equilibration by vapour diffusion, there is usually a high percentage of drops that are left clear, with neither crystals nor precipitation. There are a number of reasons why drops remain clear. One is due to the fact that the concentration of protein or precipitant, or both, is not high enough to reach supersaturation and consequently nucleation. The growth of crystals from clear drops after weeks, months or even years after the plates had been set up is likely due either to water evaporation from the reservoir through a small leak in the sealing tape (or greased cover slip), and/or proteolysis. KF13 dehydration can be employed to mimic while speeding up this slow evaporation process, thus allowing to discover new crystal hits in screening plates that had already reached vapour diffusion equilibration. To investigate this possibility, 96-well crystallization plates for lysozyme and thaumatin were prepared where some of the drops contained too low a concentration of either protein or precipitant (or both) to reach supersaturation. After 10 days, when crystal growth had not been observed for several days, fixed amounts of KF13 (3 μl) were gradually added every 3 days to the reservoirs using a multi-channel pipette. Table 4 A and B summarizes how increasing amounts of KF13 were required for crystal growth in drops with decreasing amounts of protein or precipitant (or both). Notably, the optimized crystallization conditions used to prepared lysozyme and thaumatin crystals for the KF13 cryoprotection studies discussed in previous sections contained a miminum of 0.7 M NaCl and 0.6 M NaK tartrate, respectively, as precipitants; drop dehydration via KF13 of pre-equilibrated crystallization plates produced crystal hits in drops containing as little as 0.1 NaCl and 0.05 NaK tartrate (lower precipitant concentrations than these were not tested) (Table 4A and 4B). As a negative control, the sealing tape was removed from duplicate plates, which were then left open for the same length of time that was required to add KF13 and reseal the original plates; no further crystal growth was observed in the control plates beyond that observed within the first 10 days of vapour diffusion equilibration, proving that the new crystal hits were indeed the result of KF13 drop dehydration rather than an effect of water evaporation due to the removing of the sealing tape for about 30 seconds (which is roughly the time required to add KF13 to all the reservoirs in a 96-well plate using a multi-channel pipette). Both lysozyme and thaumatin samples used in the above experiments were prepared by dissolving lyophilized protein into simple water; instead, protein samples used in crystallization typically contain salts, buffers and perhaps other chemicals (e.g., detergents in the case of membrane proteins), whose concentrations will also increase upon drop dehydration, with unpredictable effects on nucleation. This also highlights that the procedure of KF13 dehydration of clear drops is different from that of setting up new crystallization plates using a protein sample of higher concentration compared to the one used in the original plates. Another important aspect that needed to be assessed was whether KF13 dehydration of crystallization screening drops may cause the appearance of a high number of false positives due to crystallization of different salts present in most of the crystallization conditions. To investigate this, the two popular Index (Qiagen) and JCSG+ (Hampton) sparse-matrix crystallization screens were set up using a protein-less sample containing only 10 mM TrisHCl pH 8 and 300 mM NaCl (200+200 nl sitting drops and 80 μl of reservoir); the eight 96-well plates (four for each screen) were left to reach equilibrium by vapour diffusion for one week. Subsequently these plates were unsealed and 5, 10, 15 and 20 μl of KF13 were quickly added to the reservoirs of each screen, respectively, and resealed. Inspection over the next 7 days showed that none of the drops contained any crystals, either from salt or other compounds present in the many different drop conditions. This proves that KF13 dehydration is unlikely to cause an issue with false positives in hit screening experiments and at the same time it also suggests that proteins appear to play a crucial role in nucleation and subsequent crystallization of salts, which is in agreement with the fact that salt crystals are commonly obtained from different crystallization conditions depending on a specific protein sample used in the screen.

**Table 4.**
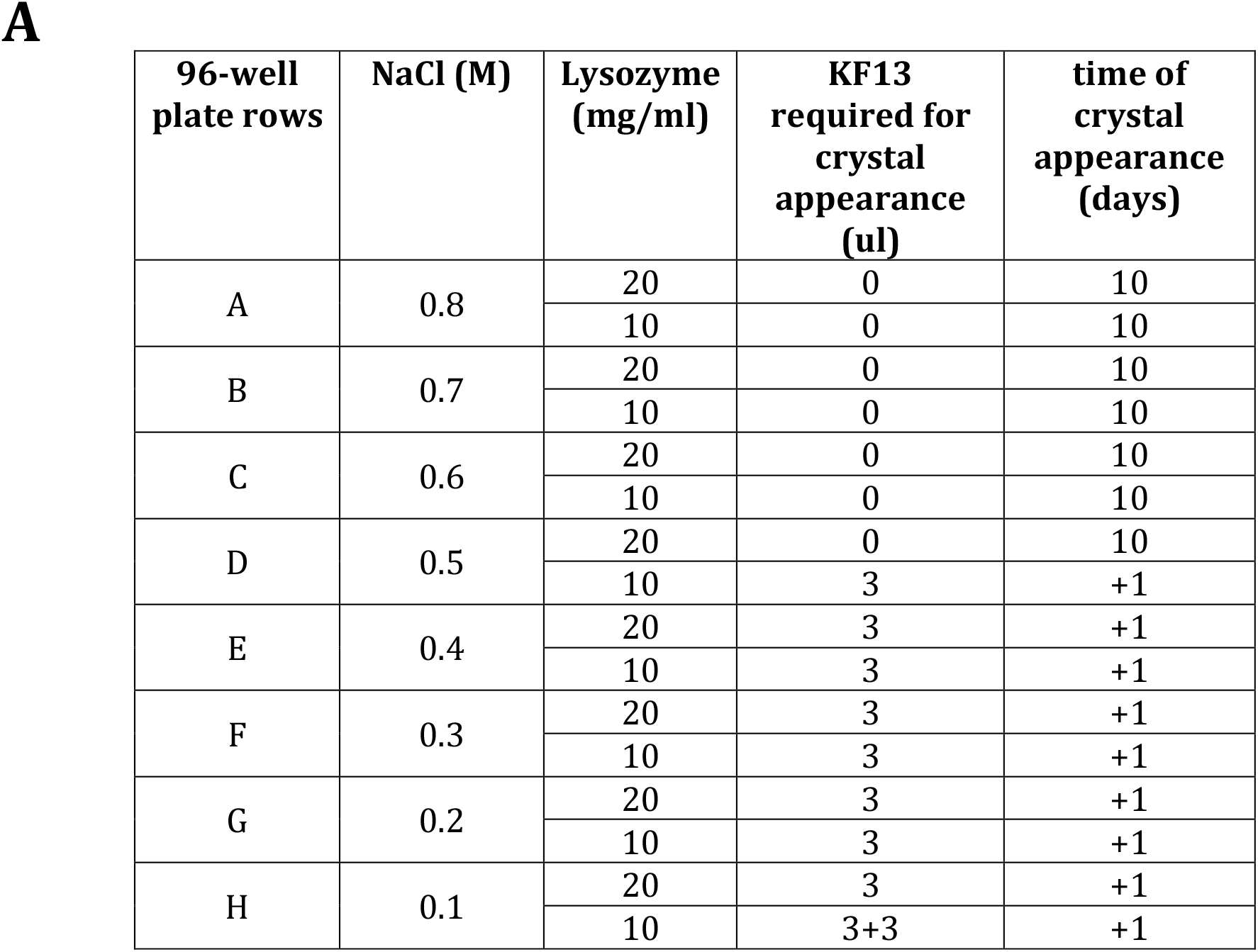
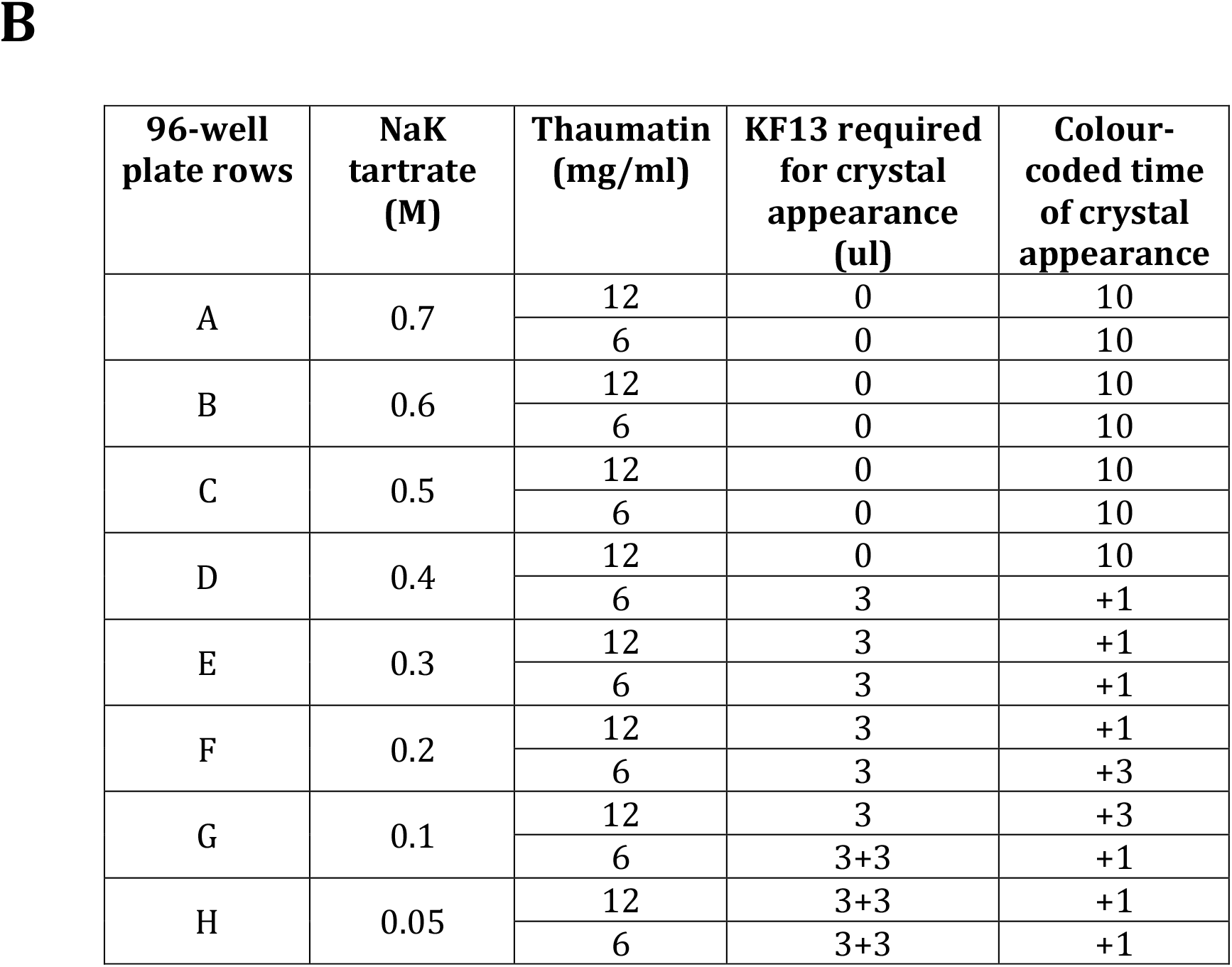
Summary of KF13 volumes required to be added to pre-equilibrated crystallisation plates to initiate crystal nucleation in clear drops of lysozyme (A) and thaumatin (B) with different precipitant concentrations. Crystallisation plates were left to equilibrate for 10 days before 3 ul of KF13 were added at intervals of 3 days; plates were monitored daily after the first 10 days of equilibration without adding any KF13.

## 4. Conclusions

This work shows that six different crystal systems were successfully cryoprotected by dehydration of the crystal drop via vapour diffusion. This was achieved by adding a very highly concentrated salt solution directly to the reservoir, without the need to manipulate the crystal drop. Screening of various salt solutions showed that 13 M potassium formate (KF13) possesses the ideal osmotic strength compared to other tested candidates to create a one-step cryoprotection protocol, since small amounts of KF13 between only 4-20% of the final volume (reservoir plus KF13) sufficed to achieve complete cryoprotection in all six crystal forms tested. Being able to limit the added KF13 volume within this range is very important because it means that a reservoir volume of up to 80% of the well capacity can be used in crystallization experiments without the need to remove any of the reservoir to achieve cryoprotection by directly adding KF13 in a single step. Being able to add KF13 without the need of removing any of the reservoir means that the crystallization plate has to remain unsealed for shorter times (minimizing crystal drop evaporation and consequently having more control over the experiment outcome and reproducibility) and the all procedures are faster to complete. Although a one-step cryoprotection protocol could also be developed using osmotically weaker solutions than KF13 by having smaller volumes of reservoir in the well allowing addition of larger volumes of the dehydrating solution, it is preferable to keep the volume of reservoir to a reasonable level (e.g., around 60-80 ul in 96-well MRC plate from SWISS-CI), since too little reservoir in the well can dry out too fast during storage (e.g., pre-filled plates stored at 10° in our crystallization facility) or even during crystallization screening experiments at room temperature. As summarized in Table 5, the KF13 protocol for protein crystal cryoprotection possesses all the qualities of the ideal method, such as being high throughput (unlike protocols for crystal freezing under high pressure or vapour diffusion of volatile alcohols), non-labour intensive (unlike the use of cryoproctective agents in a drug discovery screening experiment, where a different cryosolution would need to be prepared for each different compound/ligand), non-invasive (no crystal handling during transfer into new cryosolution drops) and not causing drop dilution by adding cryoprotective agents using an acoustic dispenser (in the case of bound ligands, this weakens binding affinities). The method described by Douangamath *et al*. (2013), which is carried out in two steps, by first completely removing the crystallization reservoir in the well and then replacing this with a solution containing NaCl, scores similarly to the KF13 protocol (Table 5). Nevertheless, while this NaCl method may be suitable for small scale projects where the manual procedure can be carried out by making a small incision in the sealing tape to minimize evaporation while executing the two steps, in high throughput cases instead, where the sealing tape is completely removed to gain easy access to all 96 wells, crystal drops remain exposed to evaporation considerably longer than during a single step procedure; resealing the plate quickly is key to achieve a slow and controlled dehydration. Moreover, completely replacing the reservoirs with a pure solution of NaCl cannot guarantee a gentle and slow dehydration rate as if gradually adding small volumes of KF13, with the latter process more likely to benefit the diffraction quality.

**Table 5.** Summary of advantages and disadvantages of cryoprotection methods available in macromolecular crystallography.

| Cryoprotection strategies | No crystal handling | No osmotic shock | No crystal drop dilution | No preparation of cryosolution with or without ligand(s) | Increase in ligand affinity by dehydration | Possible increase in diffraction resolution by dehydration | Single step protocol | High throughput |
| --- | --- | --- | --- | --- | --- | --- | --- | --- |
| KF13 | ✓ | ✓ | ✓ | ✓ | ✓ | ✓ | ✓ | ✓ |
| Crystal soaking in cryosolutions | ✗ | ✗ | ✓ | ✗ | ✗ | ✗ | ✓ | ✗ |
| Humidity control device | ✗ | ✓ | ✓ | ✓ | ✓ | ✓ | ✗ | ✗ |
| Vapour diffusion of volatile alcohols | ✗ | ✗ | ✗ | ✓ | ✗ | ✗ | ✓ | ✗ |
| Acoustic dispenser | ✓ | ✗ | ✗ | ✓ | ✗ | ✗ | ✓ | ✓ |
| Increase in environmental pressure | ✗ | ✓ | ✓ | ✓ | ✗ | ✗ | ✓ | ✗ |
| Reservoir replacement by NaCl | ✓ | ✓ | ✓ | ✓ | ✓ | ✓ | ✗ | ✓/✗ |

Cryoprotection by KF13 is very useful for drug discovery projects that are characterized by high crystal form redundancy (e.g., the amount of KF13 to be added to the reservoir to achieve optimal cryoprotection only needs to be established once) and many different ligands (which do not need to be added to KF13 since this is added to the reservoir rather than the drops); also dehydration of drops in KF13 cryoprotection can improve the ligand occupancy and/or diffraction resolution.

Moreover, this study showed that crystallization drops in pre-equilibrated plates potentially capable of producing crystals but containing too little of either protein or precipitant (or both), can be pushed into supersaturation, nucleation and crystal growth by exerting further vapour diffusion dehydration by adding KF13 to the reservoir. This aspect has application in crystallization screening experiments (both small scale and high throughput) where “idled” clear drops in plates that have already reached equilibration can be rapidly tested for being undersaturated by adding KF13 to the reservoir.

## ABBREVIATIONS

KF13: 13 M potassium formate solution
(MRC) LMB: MRC Laboratory of Molecular Biology

## Supporting information

Figures and tables showing diffraction images and crystallization plate set ups, respectively, discussed in the text.

## Acknowledgements

I thank Fabrice Gorrec and David Barford (MRC LMB) for discussions and critical reading of the manuscript; Danguole Ciziene, Christina Heroven and Stan Yatskevich (MRC LMB) for kindly donating crystals of *Staphylococcus aureus* FtsA filaments, glutamate receptor ligand binding domain in complex with agonist (GluLBD) and *Homo sapiens* hetero-pentamer Cenp-OPQUR complex, respectively.

## References

Abergel, C. (2004). Acta Crystallogr D Struct Biol 60, 1413–1416.

Berejnov, V., Husseini, N. S., Alsaied, O. A. & Thorne, R. E. (2006). Journal of Applied Crystallography 39, 244–251.

Bujacz, G., Wrzesniewska, B. & Bujacz, A. (2010). Acta Crystallogr D Biol Crystallogr. 66, 789–796.

Burkhardt, A., Warmer, M., Panneerselvam, S., Wagner, A., Zouni, A., Glockner, C., Reimer, R., Hohenberg, H. & Meents, A. (2012). Acta Crystallogr Sect F Struct Biol Cryst Commun 68, 495–500.

Collins, P. M., Ng, J. T., Talon, R., Nekrosiute, K., Krojer, T., Douangamath, A., Brandao-Neto, J., Wright, N., Pearce, N. M. & von Delft, F. (2017). Acta Crystallogr D Struct Biol 73, 246–255.

Douangamath, A., Aller, P., Lukacik, P., Sanchez-Weatherby, J., Moraes, I. & Brandao-Neto, J. (2013). Acta Crystallogr D Biol Crystallogr 69, 920–923.

Esnouf, R. M., Ren, J., Garman, E. F., Somers, D. O., Ross, C. K., Jones, E. Y., Stammers, D. K. & Stuart, D. I. (1998). Acta Crystallogr D Biol Crystallogr 54, 938–953.

Farley, C. & Juers, D. H. (2014). J Struct Biol 188, 102–106.

Fuentes-Landete, V., Mitterdorfer, C., Handle, P. H., Ruiz, G. N., Fuhrmann, S. & Loerting, T. (2015). Fuentes-Landete, V., Mitterdorfer, C., Handle, P. H., Ruiz, G. N., Fuhrmann, S. & Loerting, T.

Gulick, A. M., Horswill, A. R., Thoden, J. B., Escalante-Semerena, J. C. & Rayment, I. (2002). Acta Crystallogr D Biol Crystallogr 58, 306–309.

Haas, D. J. & Rossmann, M. G. (1970). Acta Crystallogr B. 26, 998–1004.

Heras, B., Edeling, M. A., Byriel, K. A., Jones, A., Raina, S. & Martin, J. L. (2003). Structure 11, 139–145.

Holyoak, T., Fenn, T. D., Wilson, M. A., Moulin, A. G., Ringe, D. & Petsko, G. A. (2003). Acta Crystallogr D Biol Crystallogr 59, 2356–2358.

Hope, H. (1988). Acta Crystallogr B 44 (Pt 1), 22–26.

Juers, D. H. & Matthews, B. W. (2001). J Mol Biol 311, 851–862.

Juers, D. H. & Matthews, B. W. (2004). Q Rev Biophys 37, 105–119.

Kiefersauer, R., Than, M. E., Dobbek, H., Gremer, L., Melero, M., Strobl, S., Dias, J. M., Soulimane, T. & Huber, R. (2000). J. Appl. Cryst. 33, 1223–1230.

Kim, C. U., Kapfer, R. & Gruner, S. M. (2005). Acta Crystallogr D Biol Crystallogr. 61, 881–890.

Kmetko, J., Husseini, N. S., Naides, M., Kalinin, Y. & Thorne, R. E. (2006). Acta Crystallogr D Biol Crystallogr., 1030–1038.

Kriminski, S., Caylor, C. L., Nonato, M. C., Finkelstein, K. D. & Thorne, R. E. (2002). Acta Crystallogr D Biol Crystallogr 58, 459–471.

Low, B. W., Chen, C. C., Berger, J. E., Singman, L. & Pletcher, J. F. (1966). Proc Natl Acad Sci U S A. 56, 1746–1750.

Marshall, H., Venkat, M., Seng, N. S., Cahn, J. & Juers, D. H. (2012). Acta Crystallogr D Biol Crystallogr 68, 69–81.

Owen, R. L., Rudino-Pinera, E. & Garman, E. F. (2006). Proc Natl Acad Sci U S A 103, 4912–4917.

Parkhurst, J. M., Thorn, A., Vollmar, M., Winter, G., Waterman, D. G., Fuentes-Montero, L., Gildea, R. J., Murshudov, G. N. & Evans, G. (2017). IUCrJ 4, 626–638.

Pemberton, T. A., Still, B. R., Christensen, E. M., Singh, H., Srivastava, D. & Tanner, J. J. (2012). Acta Crystallogr D Biol Crystallogr 68, 1010–1018.

Pflugrath, J. W. (2015). Acta Crystallogr F Struct Biol Commun 71, 622–642.

Rubinson, K. A., Ladner, J. E., Tordova, M. & Gilliland, G. L. (2000). Acta Crystallogr D Biol Crystallogr 56, 996–1001.

Sanchez-Weatherby, J., Bowler, M. W., Huet, J., Gobbo, A., Felisaz, F., Lavault, B., Moya, R., Kadlec, J., Ravelli, R. B. & Cipriani, F. (2009). Acta Crystallogr D Biol Crystallogr 65, 1237–1246.

Thomanek, U. F., Parak, F., Mossbauer, R. L., Formanek, H., Schwager, P. & Hoppe, W. (1973). Acta Cryst. A.

Thorn, A., Parkhurst, J., Emsley, P., Nicholls, R., Vollmar, M., Evans, G. & Murshudov, G. (2017). 729–737.

Vera, L. & Stura, E. A. (2014). Cryst. Growth Des. 14, 427–435.

Winter, G., Waterman, D. G., Parkhurst, J. M., Brewster, A. S., Gildea, R. J., Gerstel, M., Fuentes-Montero, L., Vollmar, M., Michels-Clark, T., Young, I. D., Sauter, N. K. & Evans, G. (2018). Acta Crystallogr D Struct Biol 74, 85–97.

