## Supplementary material for "A drug discovery-oriented non-invasive protocol for protein crystal cryoprotection by dehydration, with application for crystallization screening": pdf

### Supplementary information

**Table S1. Crystallisation conditions in the 96-well plates used to investigate promotion of crystal nucleation in already equilibrated crystals drops.** Each table correspond to a full 96-well plate, each letter correspond to a row with 12 identical repeats and two drops with different protein concentrations were set up for each condition.

| 96-well plate rows | Lysozyme (mg/ml) | NaCl (M) |
|---|---|---|
| A | 20 | 0.8 |
| | 10 | |
| B | 20 | 0.7 |
| | 10 | |
| C | 20 | 0.6 |
| | 10 | |
| D | 20 | 0.5 |
| | 10 | |
| E | 20 | 0.4 |
| | 10 | |
| F | 20 | 0.3 |
| | 10 | |
| G | 20 | 0.2 |
| | 10 | |
| H | 20 | 0.1 |
| | 10 | |

| 96-well plate rows | Thaumatin (mg/ml) | NaK tartrate (M) |
|---|---|---|
| A | 12 | 0.7 |
| | 6 | |
| B | 12 | 0.6 |
| | 6 | |
| C | 12 | 0.5 |
| | 6 | |
| D | 12 | 0.4 |
| | 6 | |
| E | 12 | 0.3 |
| | 6 | |
| F | 12 | 0.2 |
| | 6 | |
| G | 12 | 0.1 |
| | 6 | |
| H | 12 | 0.05 |
| | 6 | |

| 96-well plate rows | ConcanavalinA (mg/ml) | PEG 6k (%) |
|---|---|---|
| A | 10 | 10 |
| | 5 | |
| B | 10 | 9 |
| | 5 | |
| C | 10 | 8 |
| | 5 | |
| D | 10 | 7 |
| | 5 | |
| E | 10 | 6 |
| | 5 | |
| F | 10 | 5 |
| | 5 | |
| G | 10 | 4 |
| | 5 | |
| H | 10 | 3 |
| | 5 | |

### Figures

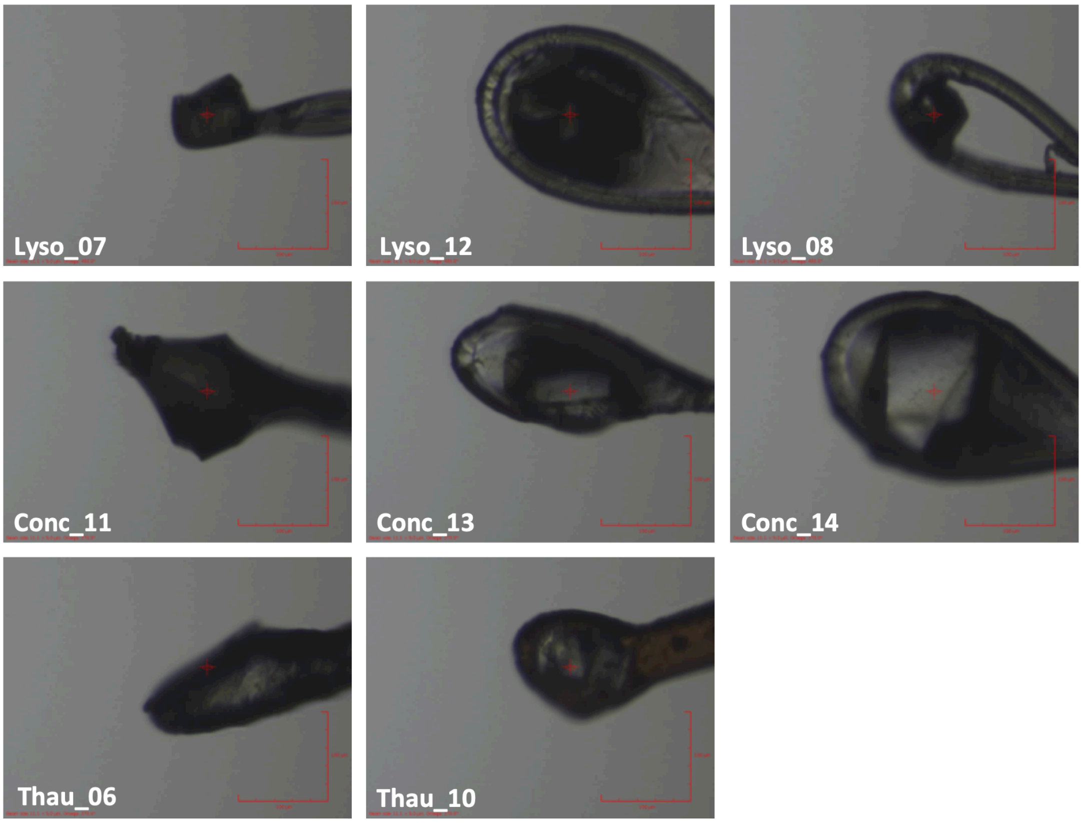


**Figure S1. Examples of choices of loop size in comparison to the size of the crystals.** Also the X-ray beam size is shown by the red circle with a cross-hair in the middle.

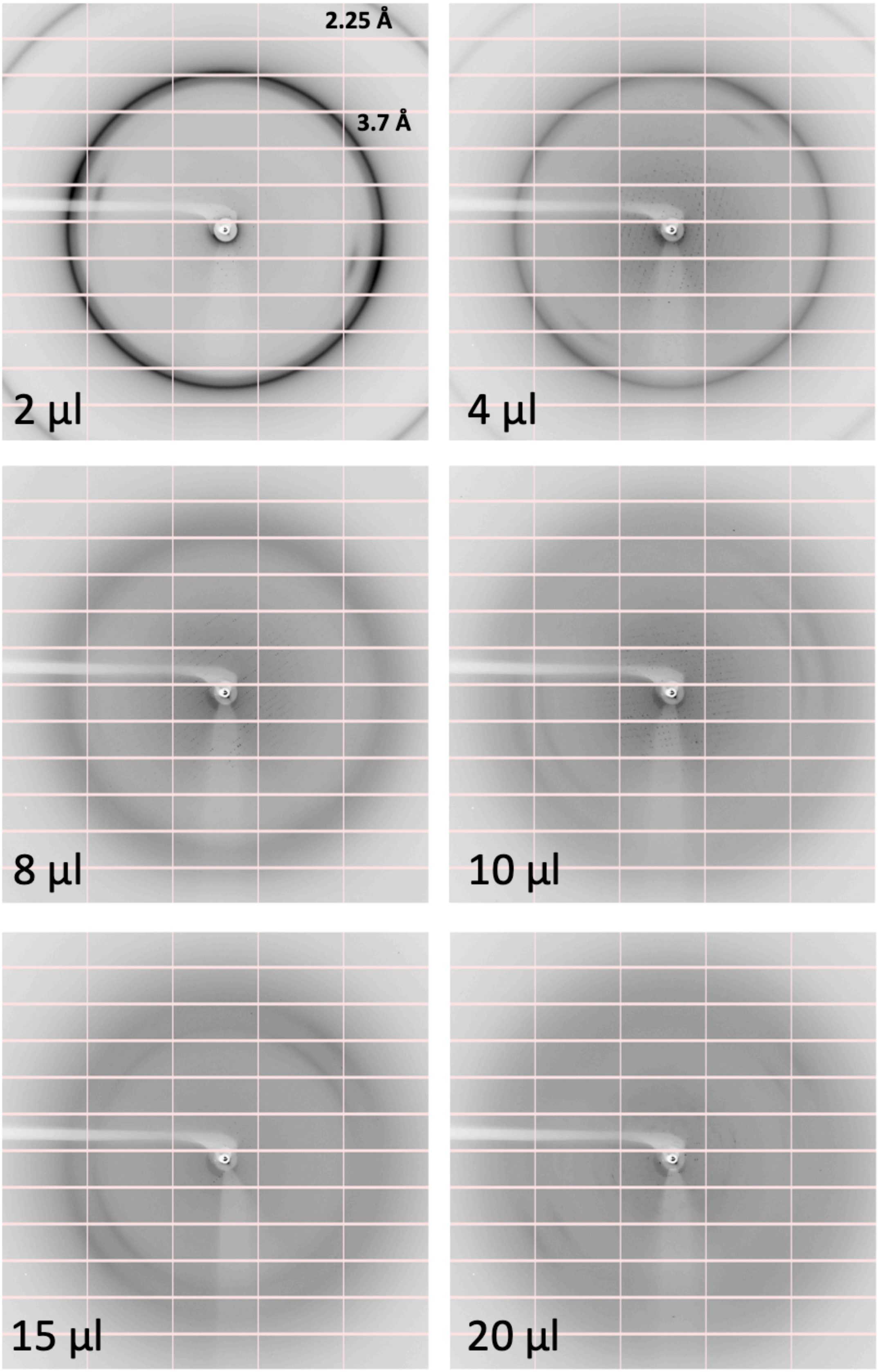


**Figure S2. Diffraction images from crystals of FtsA filaments dehydrated by adding different amount of KF13 to the reservoir.** The volume of KF13 that was used and the diffraction resolution of the ice rings are shown.

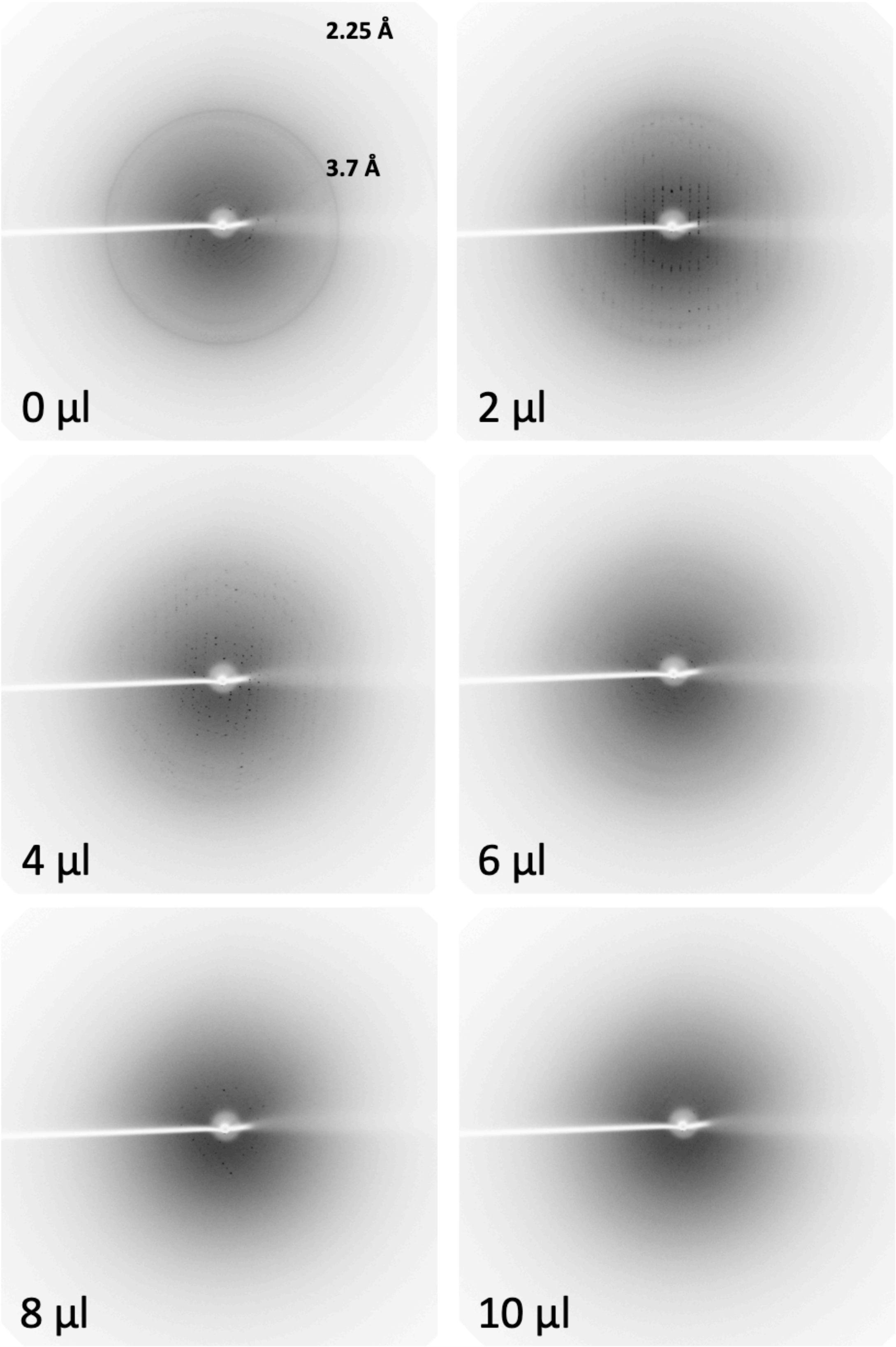


**Figure S3. Diffraction images from crystals of GluLBD in complex with agonist dehydrated by adding different amount of KF13 to the reservoir.** The volume of KF13 that was used and the diffraction resolution of the ice rings are shown.

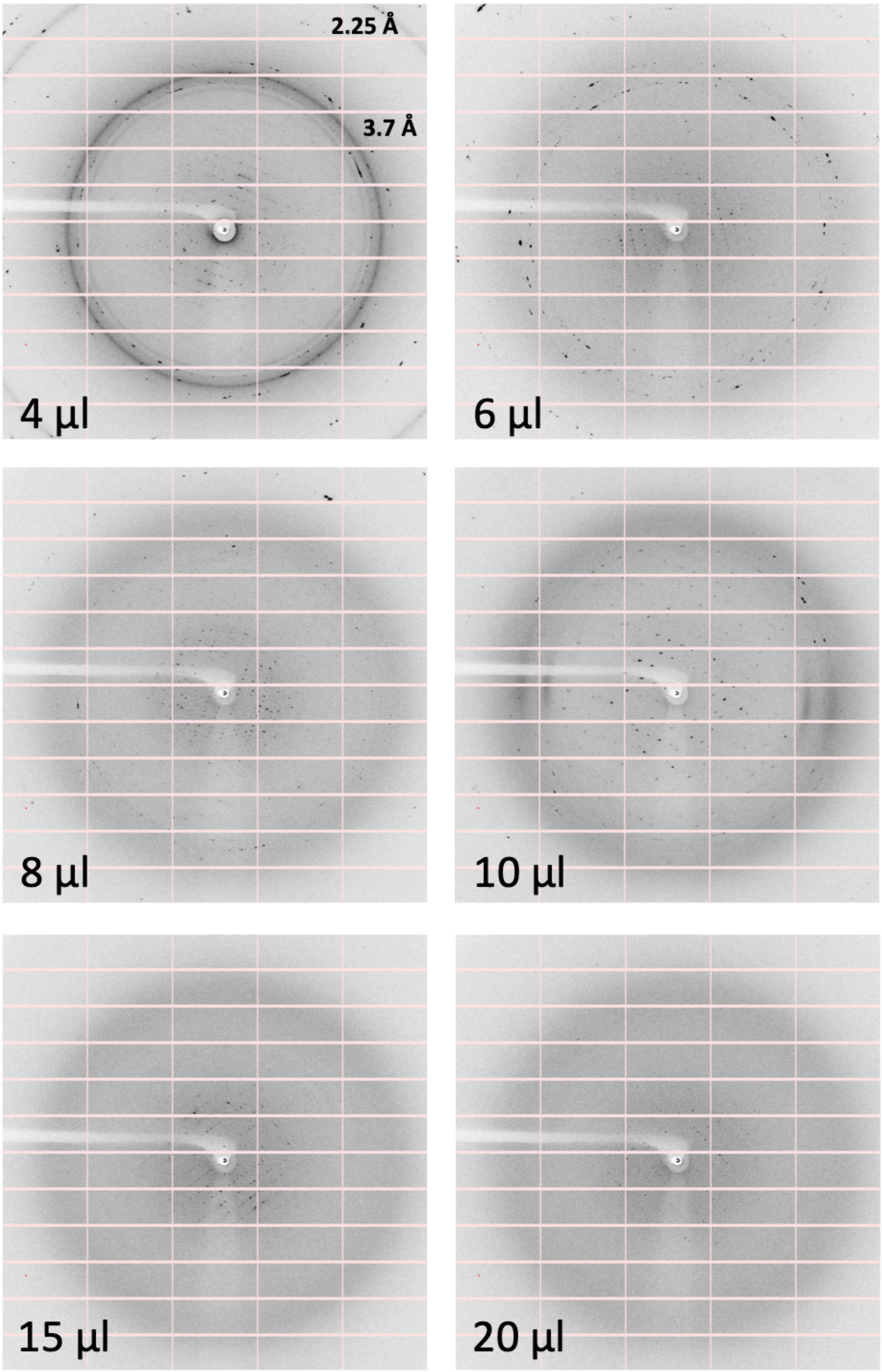


**Figure S4. Diffraction images from crystals of the hetero-pentameric complex Cenp-OPQUR dehydrated by adding different amount of KF13 to the reservoir.** The volume of KF13 that was used and the diffraction resolution

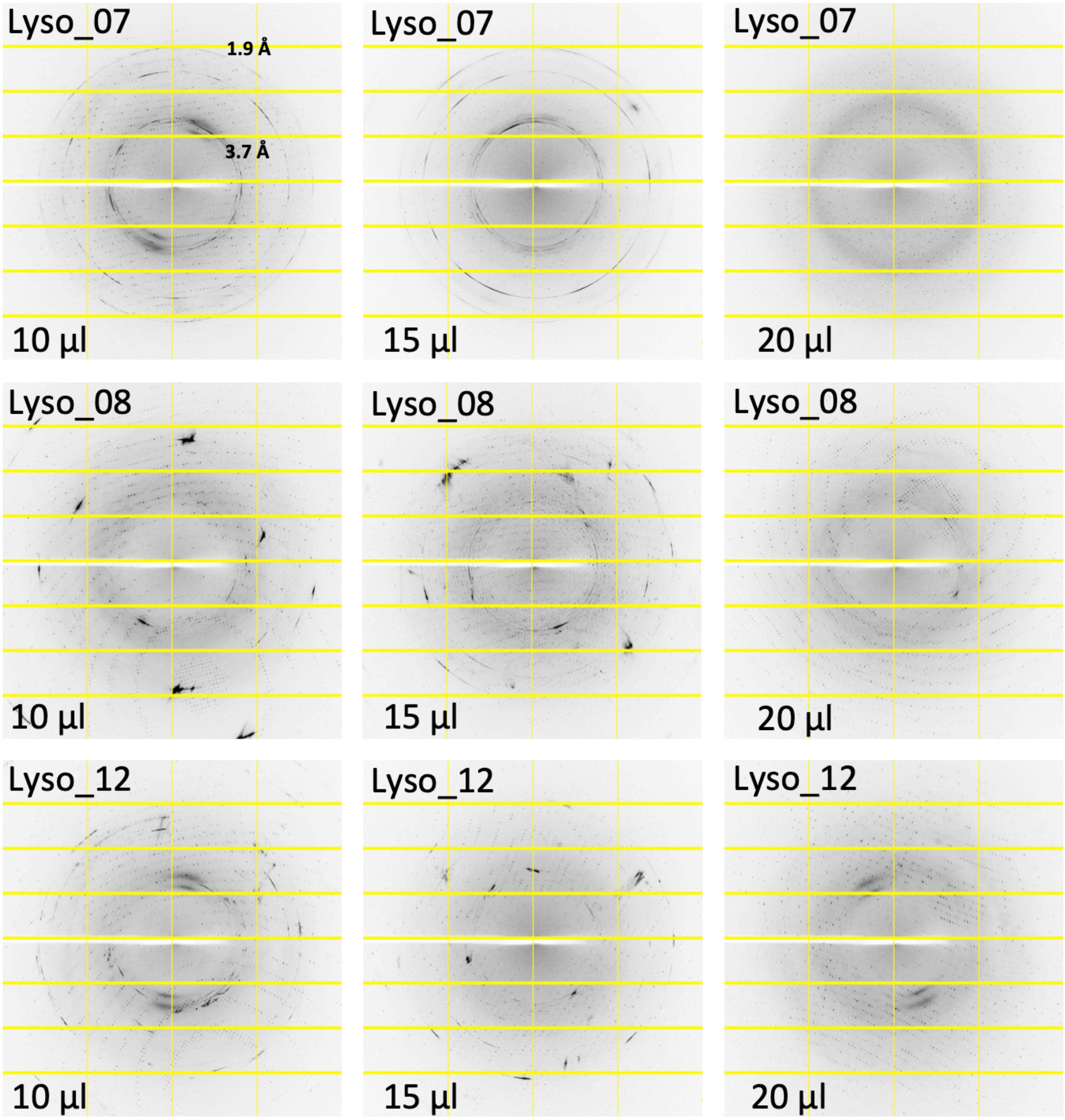

of the ice rings are shown.

**Figure S5. Diffraction images from crystals of lysozyme dehydrated by adding different amount of KF13 to the reservoir.** The volume of KF13 that was used and the diffraction resolution of the ice rings are shown. The number after the sample name indicates the amount of precipitant that was used in crystallization (see Material and Methods and Table 1).

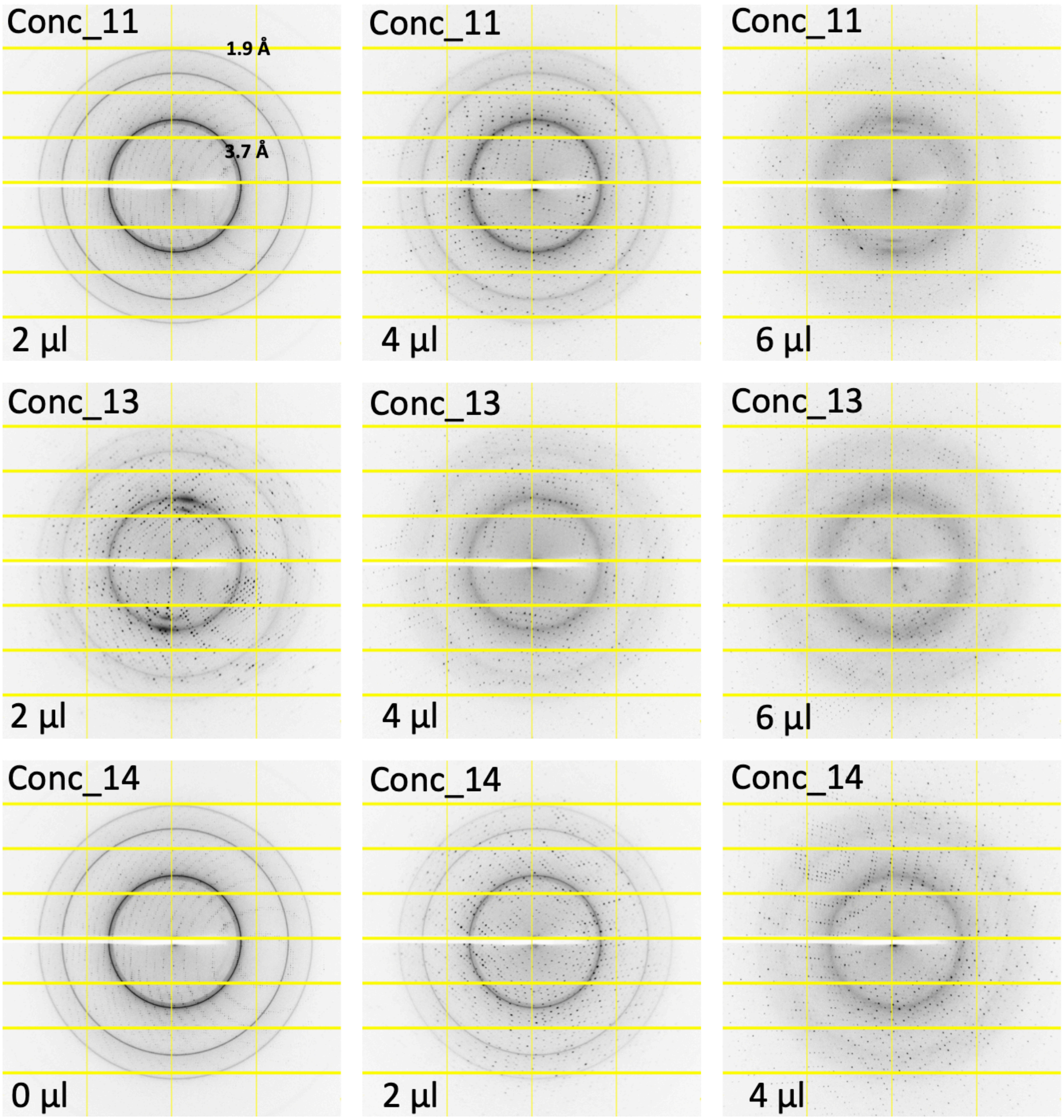


**Figure S6. Diffraction images from crystals of concanavalin A dehydrated by adding different amount of KF13 to the reservoir.** The volume of KF13 that was used and the diffraction resolution of the ice rings are shown. The number after the sample name indicates the amount of precipitant that was used in crystallization (see Material and Methods and Table 1).

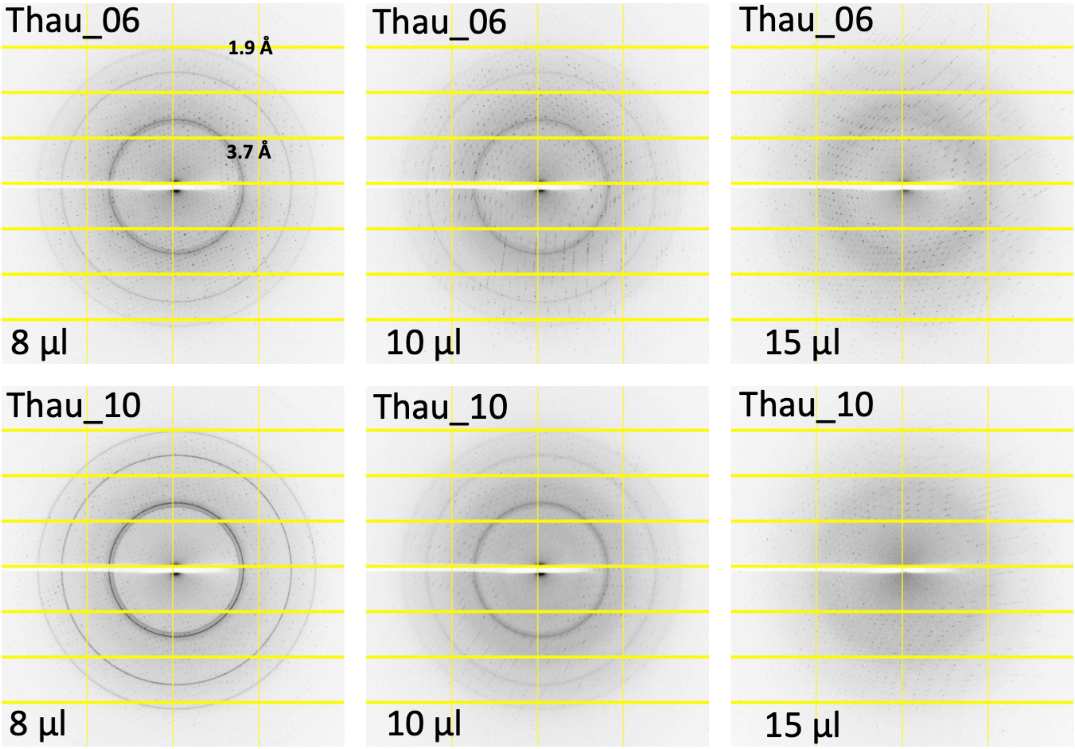


**Figure S7. Diffraction images from crystals of thaumatin dehydrated by adding different amount of KF13 to the reservoir.** The volume of KF13 that was used and the diffraction resolution of the ice rings are shown. The number after the sample name indicates the amount of precipitant that was used in crystallization (see Material and Methods and Table 1).

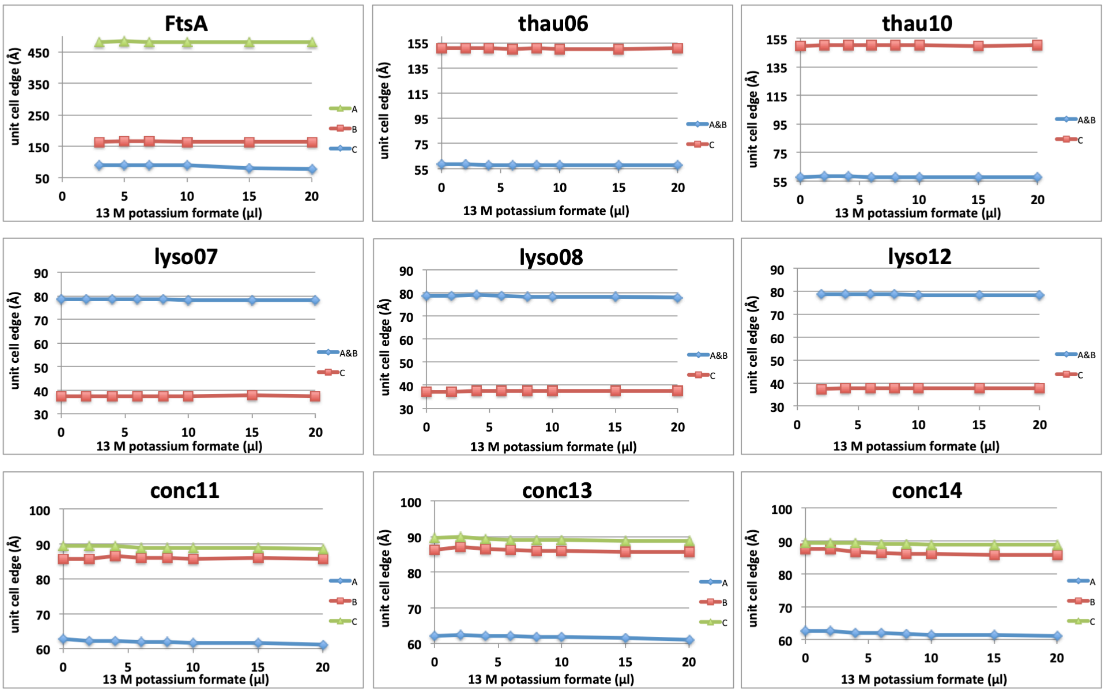


**Fig. S8. Correlation between amounts of KF13 used for crystal drop dehydration and unit cell contraction of datasets collected for different crystal samples.**
